# Towards modelling cold-water coral reef-scale crumbling: Including morphological variability in mechanical surrogate models

**DOI:** 10.1101/2022.10.06.511005

**Authors:** Marta Peña Fernández, Josh Williams, Janina V. Büscher, Jürgen Titschack, J Murray Roberts, Sebastian Henninge, Uwe Wolfram

**Affiliations:** School of Engineering and Physical Sciences, Institute of Mechanical, Process and Energy Engineering, Heriot-Watt University, Edinburgh, UK; Ryan Institute, School of Natural Sciences, University Galway, Galway, Ireland; GEOMAR Helmholtz Centre for Ocean Research Kiel, Department of Biological Oceanography, Kiel, Germany; Senckenberg am Meer, Marine Research Department, Wilhelmshaven, Germany; MARUM Center of Marine Environmental Sciences, Bremen, Germany; Changing Oceans Research Group, School of GeoSciences, University of Edinburgh, Edinburgh, UK

**Keywords:** Cold-water corals, ocean acidification, mechanical modelling, 3D morphology

## Abstract

The structural complexity of cold-water corals is threatened by ocean acidification. Increased porosity and weakening of structurally critical parts of the reef framework may lead to rapid physical collapse on an ecosystem scale, reducing their potential for biodiversity support. We can use computational models to describe the mechanisms leading to reef-crumbling. However, the implementation of such models into an efficient predictive tool that allows us to determine risk and timescales of reef collapse is missing. Here, we identified possible surrogate models to represent the branching architecture of the cold-water coral species *Lophelia pertusa*. For length scales greater than 13 cm, a continuum finite element mechanical approach can be used to analyse mechanical competence whereas at smaller length scales, mechanical surrogate models need to explicitly account for the statistical differences in the structure. We showed large morphological variations between *L. pertusa* colonies and branches, as well as *dead* and *live* skeletal structures, which need to be considered for the development of rapid monitoring tools for predicting risk of cold-water coral reefs crumbling. This will allow us to investigate timescales of changes, including the impact of exposure times to acidified waters on reef-crumbling.

## 1. Introduction

Cold-water corals (CWCs) are important ecosystem engineers, since they support high local biodiversity through the three-dimensionally (3D) complex reef habitat they make [1,2]. This structural complexity is at risk from climate-driven shifts, particularly ocean acidification, which critically affect the *dead* coral framework (i.e., erected skeleton no longer covered by soft tissue) [3–6]. Increased porosity due to ocean acidification induced dissolution in structurally critical parts of the CWC reef framework as well as the additional onset of bioerosion resulting in loss of material [7] lead to structural weakening and may result in a rapid physical habitat collapse on an ecosystem scale [4,5], thus, reducing the potential for biodiversity support [8].

Wolfram et al. [5] have recently shown that the mechanical mechanisms explaining the collapse of CWCs due to ocean acidification can be described using mathematical and computational models. The authors illustrated how changes due to ocean acidification led to a decrease of the loadbearing capacity of the skeleton using image-based finite element (FE) analysis. High-fidelity image-based models of coral structures represent a powerful tool to assess the risk of collapse of CWCs in a future ocean. However, the computational cost of this approach together with the reduced availability of 3D image data of CWCs and their reef structures restricts its use to small or only partial coral colonies and limited timepoints. To be able to estimate critical timepoints to reef-crumbling based on the time CWCs are exposed to acidified waters [4], fast and efficient surrogate models of real reef structures are essential. Such models are not existing but may, in combination with projections of seawater chemistry changes, allow us to investigate timescales of loadbearing capacity changes as well as the impact of these changes on CWC reefs overall.

Continuum FE models can provide a suitable framework to describe the local mechanical behaviour of CWC colonies using a homogenisation procedure [9,10], while reducing the computational cost of implicit simulations on large 3D image data. A continuum assumption is justified if the mechanical properties do not vary significantly over intrinsic structural dimensions [11]. As such, the effective mechanical properties of CWCs can be captured as mean values of apparent properties of statistically representative volume elements (RVEs) of the underlying structure [11,12]. These homogenised FE models require low computational resources and overcome the limitations of previous surrogate models that considered the skeletal structure uniform throughout the entire coral colony [13,14]. While the assumption of a uniform skeletal structure may hold for massive corals, it fails to account for the morphological complexity of branching corals [15–17] like the cosmopolitan scleractinian species *Lophelia pertusa* (also known as *Desmophyllum pertusum* [18]) (Figure 1).

**Figure 1.**
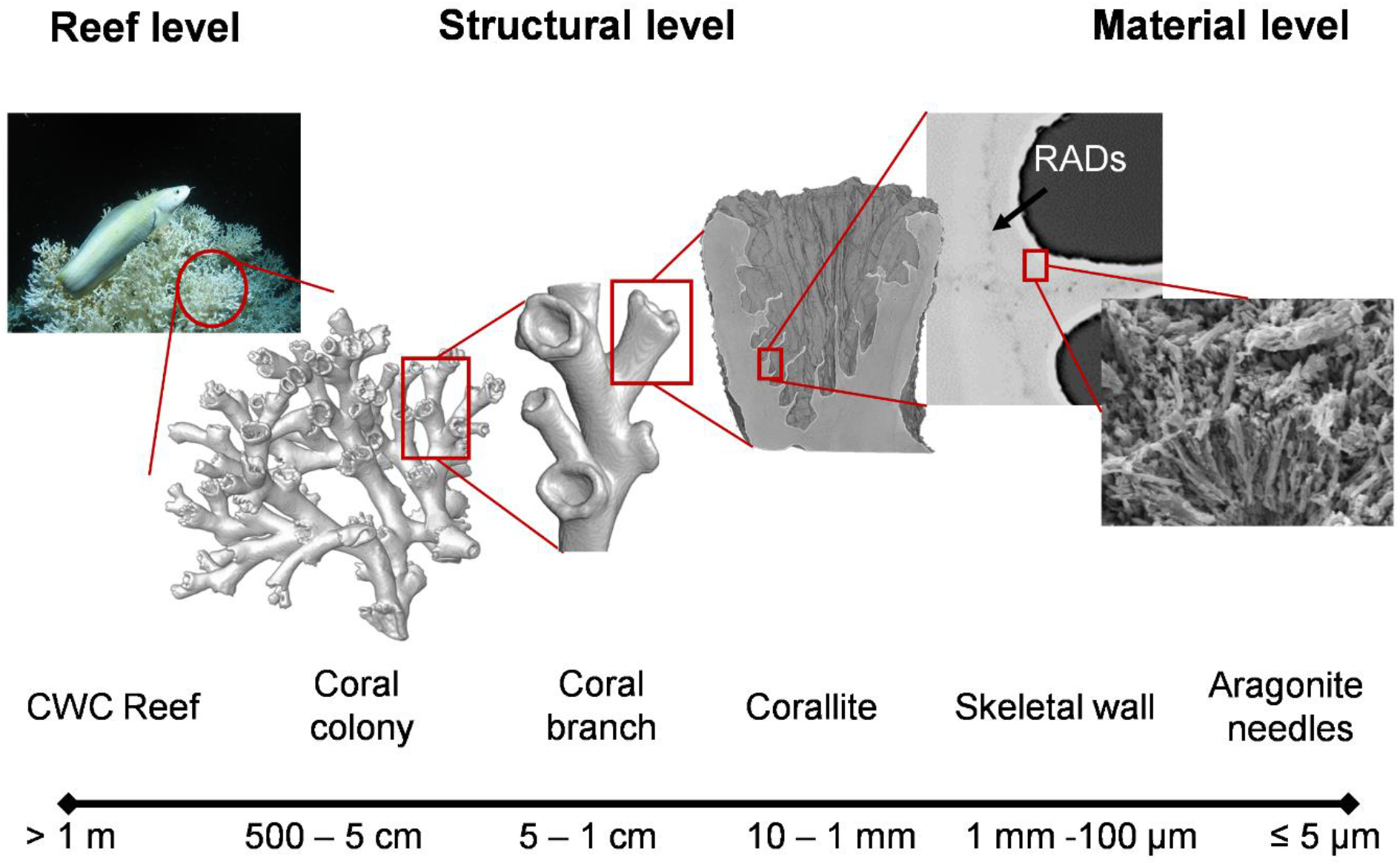
Multiscale structure of *Lophelia pertusa* cold-water coral. The material architecture of the branching cold-water coral (CWC) *L. pertusa* ranges from reef framework to aragonite crystal needles. At the material level (∼5 μm), aragonite crystals protrude from rapid accretion deposits (RADs) forming thickening bands between centres of calcification [19]. This constitutes the basic structural unit of the skeletal wall (>100 μm), which arrange into corallites at the mesoscale (∼1 cm). Corallites then assemble in a fractal-like fashion to form coral colonies, which results from the concurrent growth of multiple coral branches at the structural level [20]. These colonies form CWC reefs up to 33 m high and several kilometres in diameter [1,2,21].

To model future impacts of ocean acidification on coral skeletal integrity, it is important to understand the material architecture of CWCs (Figure 1) and the structural-mechanical relationships across length scales. While the mechanical properties at the microscale have been previously studied [4,5,22], little is known about the influence of the corallite branching arrangement on the mechanical behaviour of CWC colonies. The morphological variability of the skeletal branches needs to be considered when transitioning from structural to reef scales. Indeed, breakage of individual branches in the *L. pertusa* coral framework due to ocean acidification induced dissolution may cause the collapse of entire colonies, drastically decreasing habitat complexity. Here, we hypothesize that a critical size of a volume element for CWC skeletal structures exists, where for coral colonies larger than such critical size, a continuum FE approach can be used to investigate the risk of CWC reef collapse. Conversely, for coral colonies smaller than such critical size, the skeletal structure needs to be explicitly modelled. For both approaches, however, it is important to determine the morphological variations of CWC skeletal structure. Unlike for tropical corals that inhabit shallow waters and are easily accessible, the difficulty of collecting CWC colonies has so far restricted quantitative analysis of their morphological variations to linear measurements of small coral fragments or two-dimensional measurements from video data [20,23–26]. However, to identify critical sizes that, in turn, allow us to formulate appropriate surrogate models to capture the mechanical weakening and structural impacts of ocean acidification on exposed CWCs, an analysis of the 3D structural variations of coral skeletons across length-scales is needed. More importantly, morphological differences in *dead* and *live* CWC skeletons need to be investigated since material loss due to dissolution was observed in skeletons no longer covered by soft tissue (i.e., *dead* coral) [4].

Here, we investigate the limitations of the continuum assumption of *L. pertusa* skeletons based on the morphological variations of the skeletal structure to advance suitable surrogate models of their complex architecture. To achieve this aim, we (i) investigate the critical size of coral colonies that allow us to use a mechanical homogenisation approach to investigate crumbling and collapse of whole reef structures; (ii) analyse the morphology of *L. pertusa* coral colonies that were *alive* when collected, and *dead* erect coral framework to explain how corals occupy continuous space; (iii) characterise the branching morphology of *L. pertusa* coral skeletons to describe size and shapes of individual corallites.

## 2. Materials and methods

### 2.1. Cold-water coral specimens

We investigated morphological variations of *L. pertusa* specimens collected by Büscher et al. [27] from two Norwegian reef sites (Sula Reef Complex at 64°06.32’N, 8°07.1’E and 303 m depth, and Leksa Reef at 63°36.46’N, 9°22.76’E, 157 m depth and 63°36.43’N, 9°22.45’E, 152 m depth). *Live* coral colonies of white and orange colourmorphs as well as *dead* erect coral framework were sampled from both sites.

To investigate the continuum assumption for CWC skeletal structure, we examined two *L. pertusa* specimens collected from Rockall Bank (57°54.9’N, 13°52.296’W, unknown depth) and West Shetland (60°43.188’N, 2°55.788’W, unknown depth), which provided a larger representation of the skeletal structure (Table 1).

**Table 1.** Summary of size characteristics of the analysed CWC specimens. Volume, surface area, and dry weight are reported as median [minimum, maximum] for CWCs from Mid-Norway.

|  | Volume in cm<sup>3</sup> | Surface area in cm<sup>2</sup> | Dry weight in g |
| --- | --- | --- | --- |
| Rockall Bank (*n* = 1) | 535.0 | 649.5 | 1296.0 |
| West Shetland (*n* = 1) | 419.2 | 4204.8 | 1214.0 |
| Mid-Norway *dead* framework (*n* = 19) | 111.6 | 1632.9 | 170.3 |
|  | [52.8, 143.5] | [817.7, 1957.9] | [93.5, 226.9] |
| Mid-Norway orange corals (*n* = 11) | 15.5 | 229.6 | 28.3 |
|  | [7.0, 45.2] | [103.0, 676.8] | [13.8, 96.3] |
| Mid-Norway white corals (*n* = 11) | 28.9 | 456.6 | 59.6 |
|  | [7.8, 41.4] | [140.1, 572.3] | [13.6, 91.6] |

### 2.2. Image acquisition and processing

Computed tomography (CT) images from Büscher et al. [27] of dried coral fragments from Norwegian reefs were produced with a Toshiba Aquilion 64 clinical CT (120 kV, 600 mA, 0.351 mm in-plane pixel size, 0.5 mm slice thickness, 0.3 mm slice spacing). Images were reconstructed with a voxel size of 0.351×0.351×0.3 mm^3^. For the large specimens, we performed CT with a Siemens Somatom clinical CT (120 kV 80 mA, 0.6 mm slice thickness, 0.35 mm slice spacing). The CT images had an in-plane pixel size of 0.662 mm for the Rockall Bank and 0.445 mm for the West Shetland specimen.

We resampled the images to an isotropic voxel size (0.351 mm^3^, 0.662 mm^3^, and 0.445 mm^3^ for the Norwegian, Rockall Bank, and West of Shetland specimens, respectively) and reduced noise using a Gaussian filter. Thereafter, we segmented coral skeletons and cavities individually (Figure 2a-f) and quantified the local size and shape of both the coral skeleton and corallites (Figure 2g-f). Further details on image post processing may be found in the supplementary material section S1.

**Figure 2.**
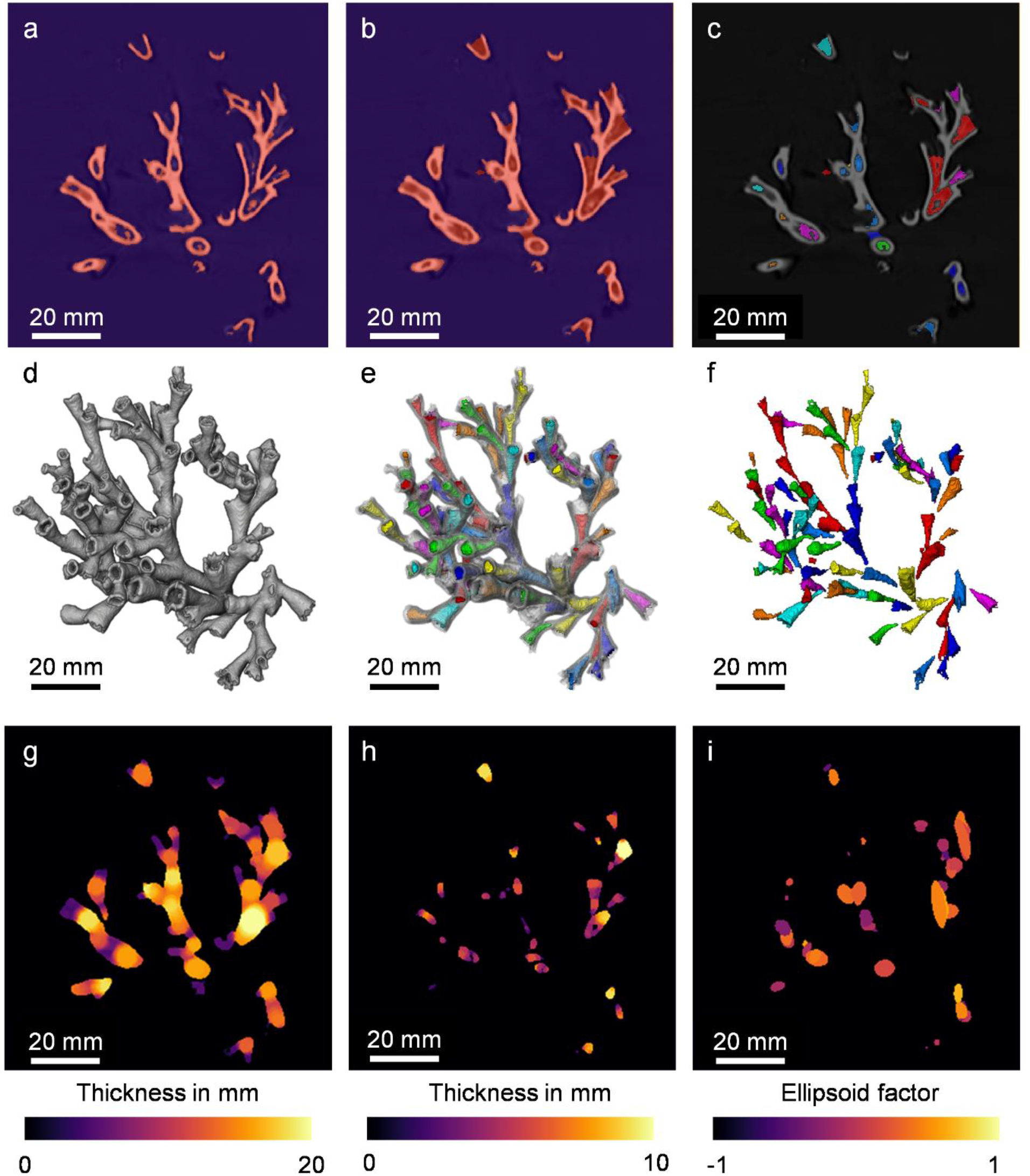
Segmentation of skeleton and cavities for an exemplary CWC specimen of the Leksa Reef. A representative CT cross-section and 3D render of the specimen are shown. (a, d) Coral skeleton was segmented. (b, e) A mask contour image was created by filling the cavities within the skeleton. (c, f) Cavities were segmented subtracting the skeleton from the masked contour and individually labelled. (g, h) The local diameter of the skeleton and branch thickness were computed from the mask contour image and the cavities image. (i) The shape of the coral branches was computed using the ellipsoid factor, where a value of −1 indicates a highly oblate shape and a value of 1 a highly prolate shape.

### 2.3. Critical size of representative volume element (RVE) of cold-water corals

We used the FE method to approximate the critical size for an RVE (supplementary material section S2) of CWC skeleton by analysing the size dependence of the elastic symmetries and properties of the structure [28]. These properties can be estimated from the stiffness tensor, 𝕊, using a direct mechanics approach through an optimisation procedure where the best orthotropic representation of 𝕊 may be found [29].

We virtually extracted the largest cuboid volume element (CVE) fully occupied by the structure (114.2 mm x 58.9 mm x 114.2 mm) from the CT reconstruction of the Rockall Bank and West Shetland specimens (Figure 3). From this, we generated 64 CVEs for each specimen with edge lengths varying from 114.5 mm to 34 mm in two directions while keeping the shorter edge constant. We created FE models by direct conversion of image voxels into isotropic linear hexahedral elements [30,31]. Additionally, we generated a mirrored model (Figure 3c) to investigate larger skeleton sizes than the ones physically available for scanning, from which we generated 217 CVEs and associated FE models with edge lengths varying from 225 mm to 45 mm, following Pahr et al. [32]. All models were analysed in Abaqus/Standard (v6 R2018, Simulia) using kinematic uniform boundary conditions, where six independent load cases (three uniform longitudinal compressive and shear strains) were applied [32]. The material was assumed to be isotropic with a Young’s modulus of 65.7 GPa and Poisson’s ratio of 0.29 [5].

**Figure 3.**
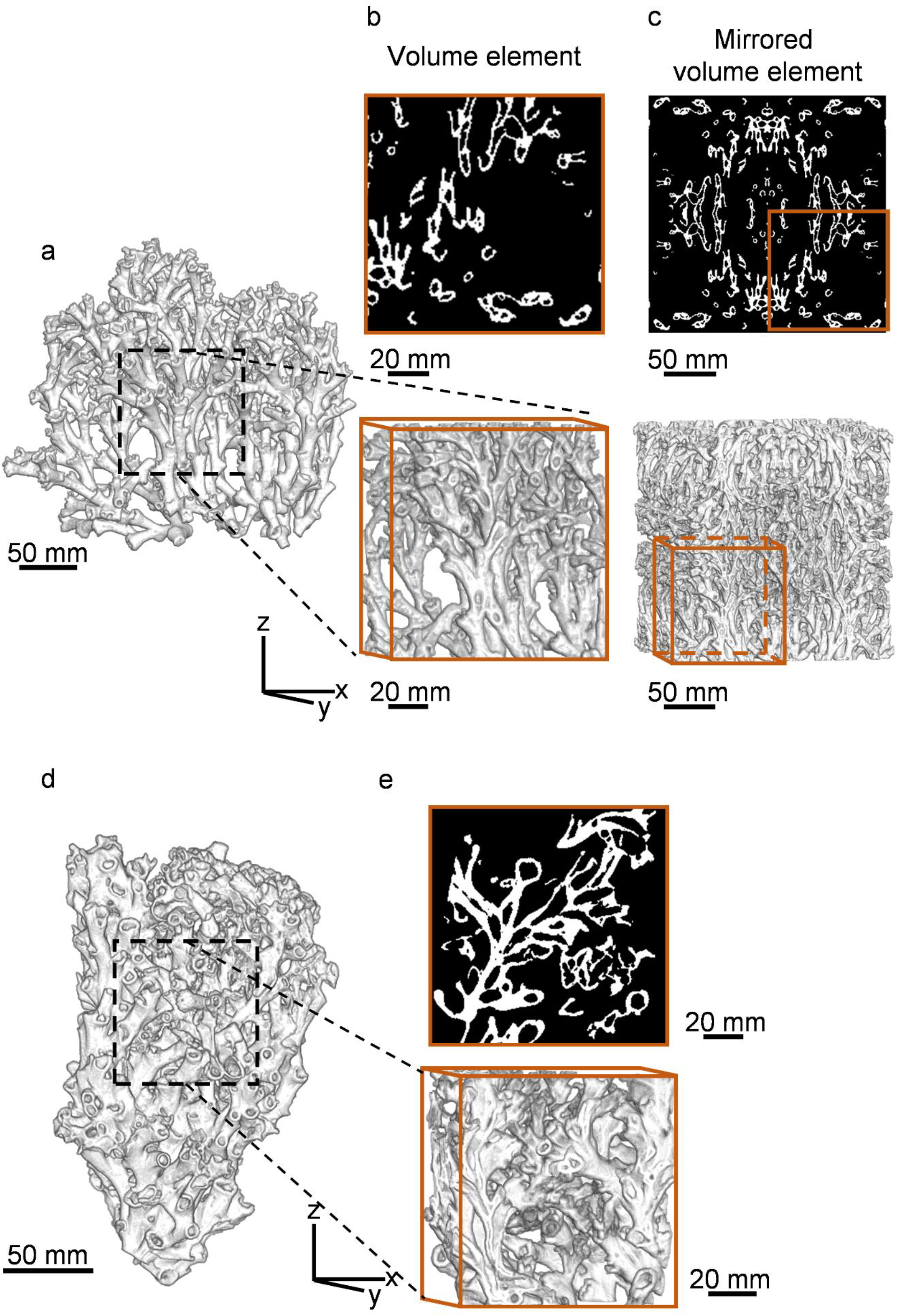
Representative volume element of large cold-water coral specimens from Rockall Bank and West Shetland. 3D render of (a) Rockall Bank and (d) West Shetland cold-water coral specimens used for finite element analysis. (b), (e) 2D cross-sections (top) and 3D volume elements (bottom) with 114.2 mm edge length in x and z axis and (c) mirrored model with 225 mm edge length.

For each model, 𝕊 was derived from the FE analysis via the apparent stresses and strains as in [32]. We calculated the orientation of the closest orthotropic stiffness tensor, 𝕊^*OPT*^, by minimising the objective function defined by:

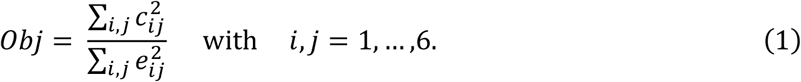

Where *c*_*ij*_ represents the nonorthotropic terms of 𝕊 and *e*_*ij*_ the orthotropic terms of the transformed 𝕊 [29]. The optimisation approach resulted in the best possible orthotropic representation of the stifness tensor, 𝕊^*OPT*^. We defined an orthotropic approximation of 𝕊^*ORT*^ by setting the nonorthotropic components to zero. The accuracy of the orthotropic assumption was quantified using the error of the orthotropic approximation [32], defined as:

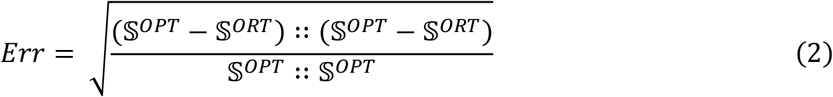

We analysed the convergence of the error with respect to the size of the CVEs as well as skeletal volume fraction (*V*_*S*_*/V*_*T*_, ratio between skeletal volume, *V*_*S*_, and total volume, *V*_*T*_). Both 𝕊^*OPT*^ and 𝕊^*ORT*^ were checked to confirm they were positive definite.

### 2.4. Morphology of CWC colony fragments

We introduce six shape variables to quantify the morphology of the CWC skeletons at the colony level based on the segmented coral skeleton images (Figure 2a, d). We quantified shape through sphericity and sparsity (capturing volume compactness), surface area to volume ratio and fractal dimension (capturing surface complexity), as well as elongation and flatness (capturing the aspect ratio of the colony) as:

i. **Sphericity** (*S*_*ph*_) is an invariant measurement of the compactness of an object’s volume. *S*_*ph*_ is defined as the ratio between the surface area of a sphere with the same skeletal volume (*V*_*S*_) as the coral skeleton and the surface area (*SA*) of the coral skeleton.

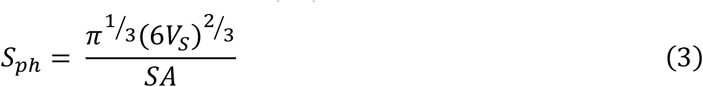
ii. **Sparsity** (*S*) is an invariant measurement of the degree to which there is space between different regions of the coral structure. *S* is defined as the ratio between the volume of an ellipsoid (*V*_*E*_) fitting the coral colony and the skeletal volume (*V*_*S*_) of the coral.

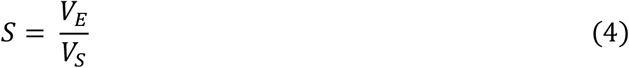
iii. **Surface area to volume ratio** (*SA*: *Vol*) refers to the amount of surface area (*SA*) per unit volume of the skeletal volume (*V*_*S*_) of the coral colony.

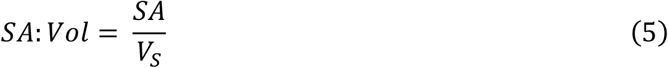
iv. **Fractal dimension** (*F*_*D*_) captures how the surface of the coral skeletal structure fills space, and it is an estimate of the spatial complexity [33]. *F*_*D*_ is computed as the slope of the number of boxes at a size *s* that contains part of the coral skeletal structure (*N*^*s*^) and the size of the boxes.

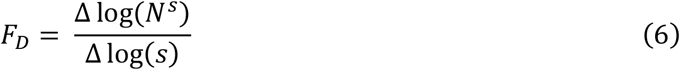
v. **Elongation** (*E*) is derived from the shape orientation of the coral colony. *E* is computed as the ratio between the length of first (*l*_1_) and second (*l*_2_) axis of an ellipsoid fitting the coral colony.

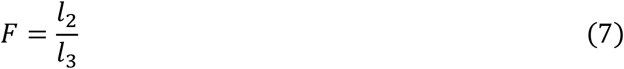
vi. **Flatness** (*F*) is the ratio between length of second (*l*_2_) and third (*l*_3_) axis of an ellipsoid fitting the coral colony.

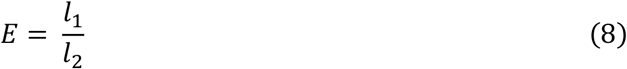

### 2.5. Morphology of CWC skeletal branches

To quantify morphological variations of the coral specimens at the branch level (i.e., size and shape of the individual and/or group of corallites), we first performed a skeletonisation [34] of the mask image contour (Figure 3b, e) via a 3D thinning algorithm and we converted the skeletonised image into a graph object (Figure 4) using the NetworkX package [35].

**Figure 4.**
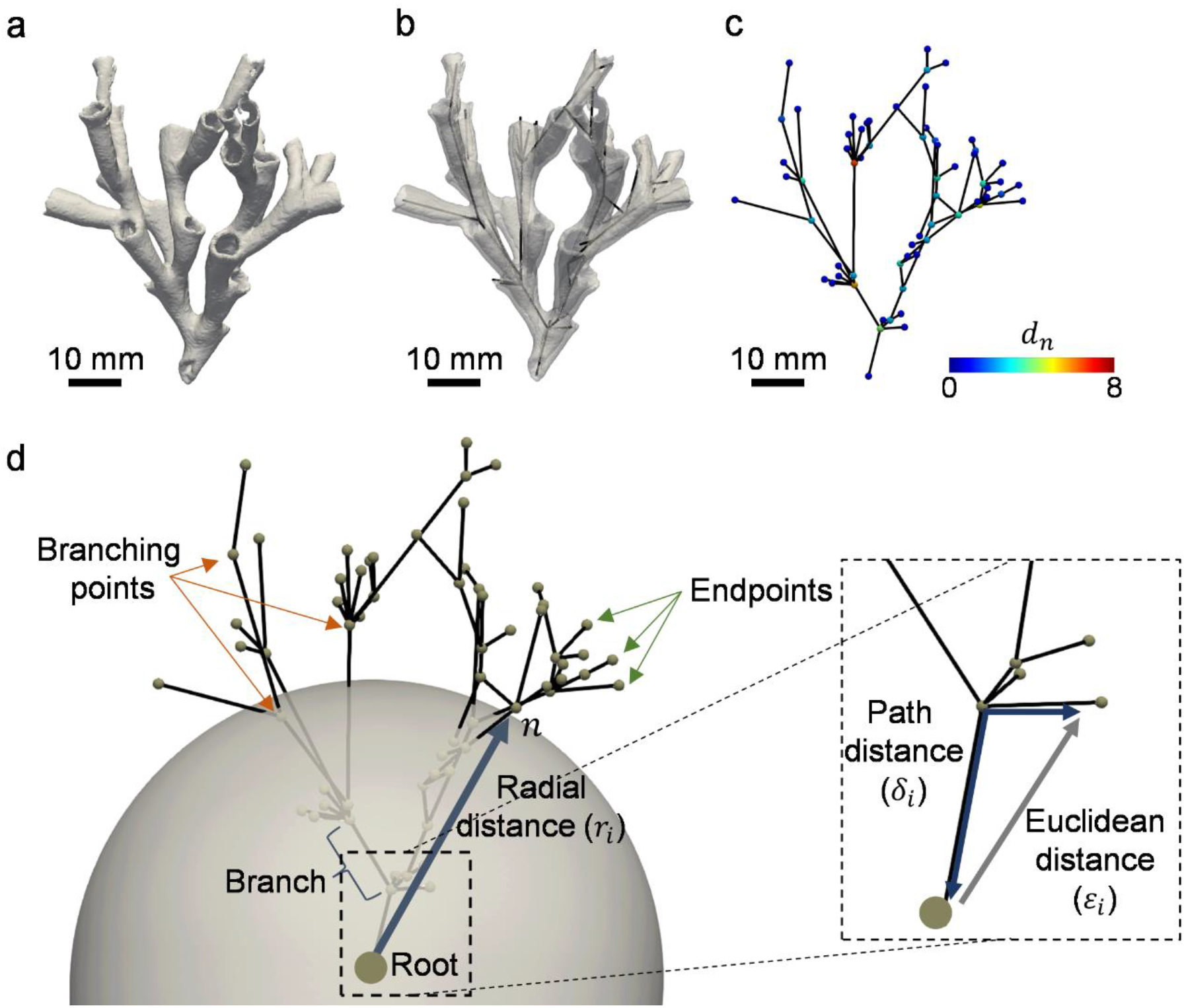
Graph representation of an exemplary CWC specimen. (a) 3D render of a coral and (b) corresponding graph representation, *G* = (*N, E, α*), in 3D space. (c) The degree of each node, *d*_*n*_, allows to classify nodes and reduce the graph to branching points (*d*_*n*_ ≥ 3) and endpoints (*d*_*n*_ = 1), by removing all path nodes (i.e., nodes within a branch, *d*_*n*_ = 2), so that edges represent the branches of the coral specimens. (d) The final coral is represented by a set of nodes, *n*, and branches (i.e., edges) with size and shape attributes, *α*, which can be analysed using topological descriptors that vary as a function of either the radial distance, *r*_*i*_, or the path distance, *δ*_*i*_ from the root.

Initially, we assigned a 3D spatial coordinate to each node based on the image coordinates and we then inspected the resulting graphs and manually selected the root (i.e., base) of each skeleton based on the morphology (Figure 4d), from where a newly oriented graph was created via a depth-first-search algorithm [36]. We added the mean coral branch thickness (Br.Th), length (Br.Len), area (Br.Ar), volume (Br.Vol), taper rate (Br.Tr), and ellipsoid factor (Br.Ef) as nodal attributes to account for the size and shape of each individual branch.

We then introduced four topological descriptors that represent morphological features of the branching coral structure, adopting those proposed by Khalil et al. [37] to study neuronal morphology. These descriptors associate a function to a given coral skeleton whose independent variable is either the path, *δ*, or radial, *r*, distance from the skeletal root (Figure 4d).

i. **Branching pattern** (*B*_*P*_) quantifies the skeletal complexity of the coral specimens and the distribution of the branches. *B*_*P*_ is related to skeletal growth and spatial arrangement and it can be defined as a function of the radial distance from the root, *r*, as:

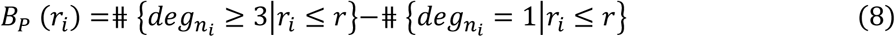

Where ⋕ represents the cardinality of each set and *r*_*i*_ the radial distance of *i*-th node, *n*_*i*_, to the root.
ii. **Terminal branch index** (*T*_*BI*_) counts the number of end points that can be reached from a given node. *T*_*BI*_ quantifies the hierarchical branching growth of CWC skeletons and it is defined as a function of the path distance, *δ*, from the root along the branches as:

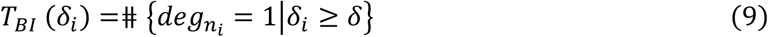

Where *δ*_*i*_ represents the path distance from the *i*-th node to the root.
iii. **Tortuosity** (*τ*) is defined as the ratio of the path distance, *δ*, by the Euclidean distance, *ε*, between a node and the root. *τ* is related to the branch growth mechanism, and it is defined as a function of the radial distance from the root, *r*, as:

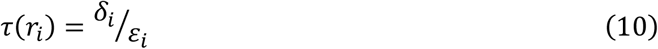

Where *ε*_*i*_ represents the Euclidean distance from the *i*-th node to the root.
iv. **Volume distribution** (*V*_*d*_) gives a measure of how the volumetric mass of the branches are distributed relative to the root. For a given coral skeleton, we consider all its nodes as cloud points in 3D space, *n*_*i*_(*x*_*i*_, *y*_*i*_, *z*_*i*_), each of them carrying a weight equal to the volume of the branch, *v*_*i*_. This volume affects the space around them such that each node contributes to a field, *V*_*i*_, which is normalised to have length *v*_*i*_ and which is of the form:

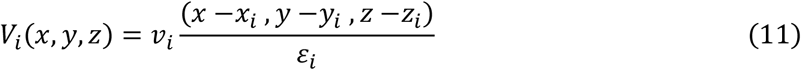

By superimposition, the node configuration of the skeleton gives rise to a vector field,*V*, whose magnitude is used as *V*_*d*_, and it is defined as a function of the radial distance to the root, *r*, as:

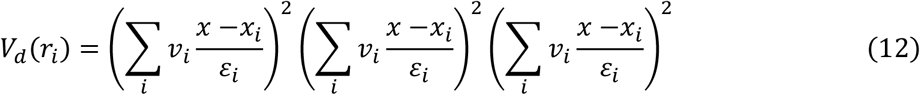

To reduce the dimensionality of the proposed descriptors as well as the coral branch attributes (i.e., Br.Th, Br.Len, Br.Ar, Br.Vol, Br.Tr and Br.Ef) we combined those measurements into a vector, *V*_*C*_, by considering the area under the curve defined by each descriptor, *ø*, as:

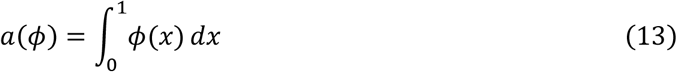

Where *x* is the normalised path, *δ*_*i*_, or radial, *r*_*i*_, distance from the skeletal root. The corresponding vector for each coral skeleton, *V*_*C*_, is:

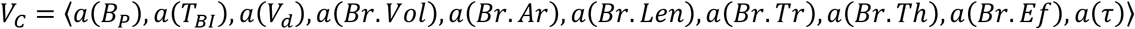

This vectorization allows us to optimize classification of the data [37].

### 2.6. Analysis of morphological parameters

Statistical analyses of morphological parameters coral skeletons were conducted in RStudio (Version 1.1.456). We used quantile–quantile plots and Shapiro–Wilk post-hoc tests to test normal distribution of data. If normality was given, we compared groups using Student’s t-tests. Where data were non-normal, we used Wilcoxon rank sum tests. We assumed a significance level of p = 0.05.

We used principal component analysis (PCA) to visualise patterns of morphological variations at the colony and branch level. We standardised variables with a mean of zero and unit variance to reduce the influence of variable scale on the projection. We used score plots to visualise the projection of each coral specimen onto the span of the two first principal components and how each group relate to each other. Finally, we investigated the relationships between the first two principal components and the original morphological variables using loading plots.

## 3. Results

### 3.1. Critical size of representative volume element of CWCs

The error of the orthotropic approximation of the stifness tensor decreases with increasing specimen size (Figure 5a). The error decreased significantly after 6 cm edge length and converged to less than 3% at ∼9 cm edge length. Considering that the Br.Sp was 2.57 cm for the Rockall Bank coral specimen and 1.75 cm for the West Shetland specimen, the error converged at 4 to 5 Br.Sp (Figure 5b). The error also decreased for increasing *V*_*S*_*/V*_*T*_ (Figure 5c). The size of the volume elements also affected the Young’s and shear modulus but had minimal influence in the Poisson’s ratio (supplementary Figure S1). Similar convergence was observed for the mirrored models (*Err* < 2.5% at ∼9 cm edge length, ∼4 Br.Sp). The imposed orthotropic structure in those models resulted in *Err* < 1.5% and lower standard deviation for extracted volume elements >13 cm. Therefore, a critical size of ∼13 cm (5 to 6 Br.Sp), should provide sufficiently average continuum quantities, thus, allowing for a mechanical homogenisation approach for *L. pertusa* skeletal structures.

**Figure 5.**
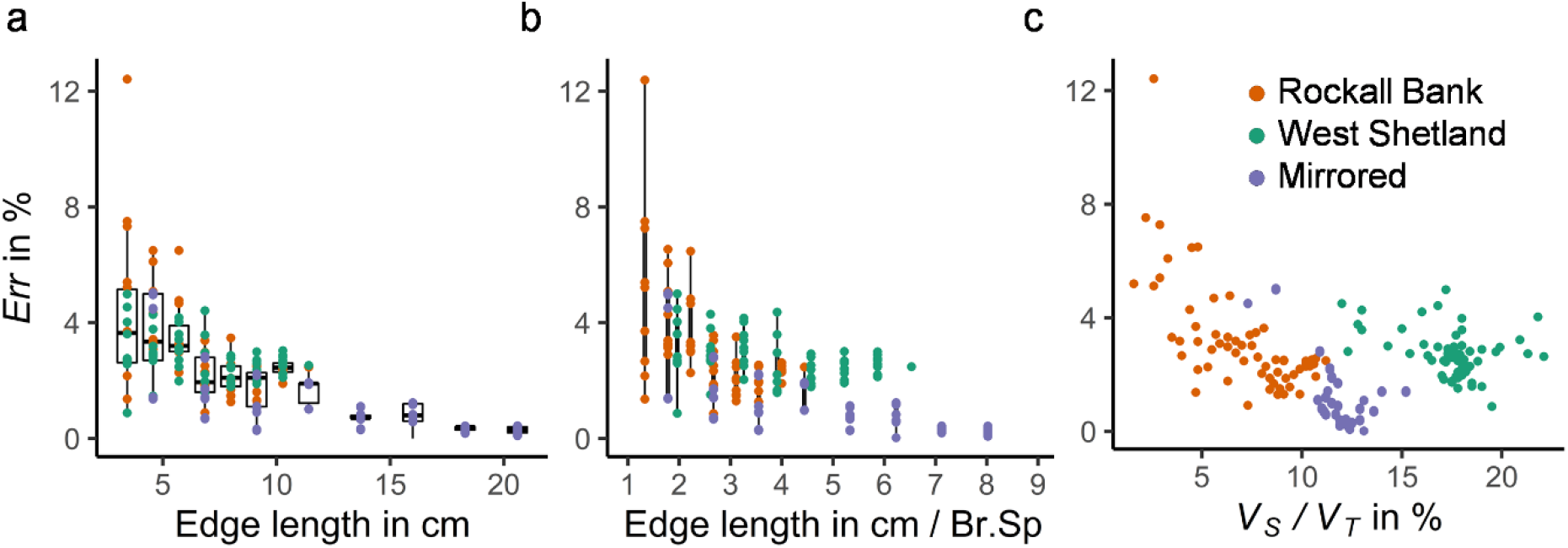
Error of orthotropic approximation. Boxplots of the errors for the analysed volume elements of the Rockall Bank, West Shetland and mirrored models as a function of the (a) edge length of the cuboid volume elements and (b) edge length expressed in number of mean branch spacing (Br.Sp) and (c) skeletal volume fraction (*V*_*S*_*/V*_*T*_).

### 3.2. Morphology of CWC colony fragments

*Live* CWC fragments exhibited significantly greater SA:Vol and a more compact structure (i.e., higher sphericity and sparsity) compared to *dead* specimens (Figure 6). The complexity of the *dead* CWC framework was demonstrated through the higher fractal dimension. *Dead* framework displayed also a more elongated shape. *Live* orange CWC specimens showed larger variability of the shape parameters, but no significant differences were observed between colourmorphs.

**Figure 6.**
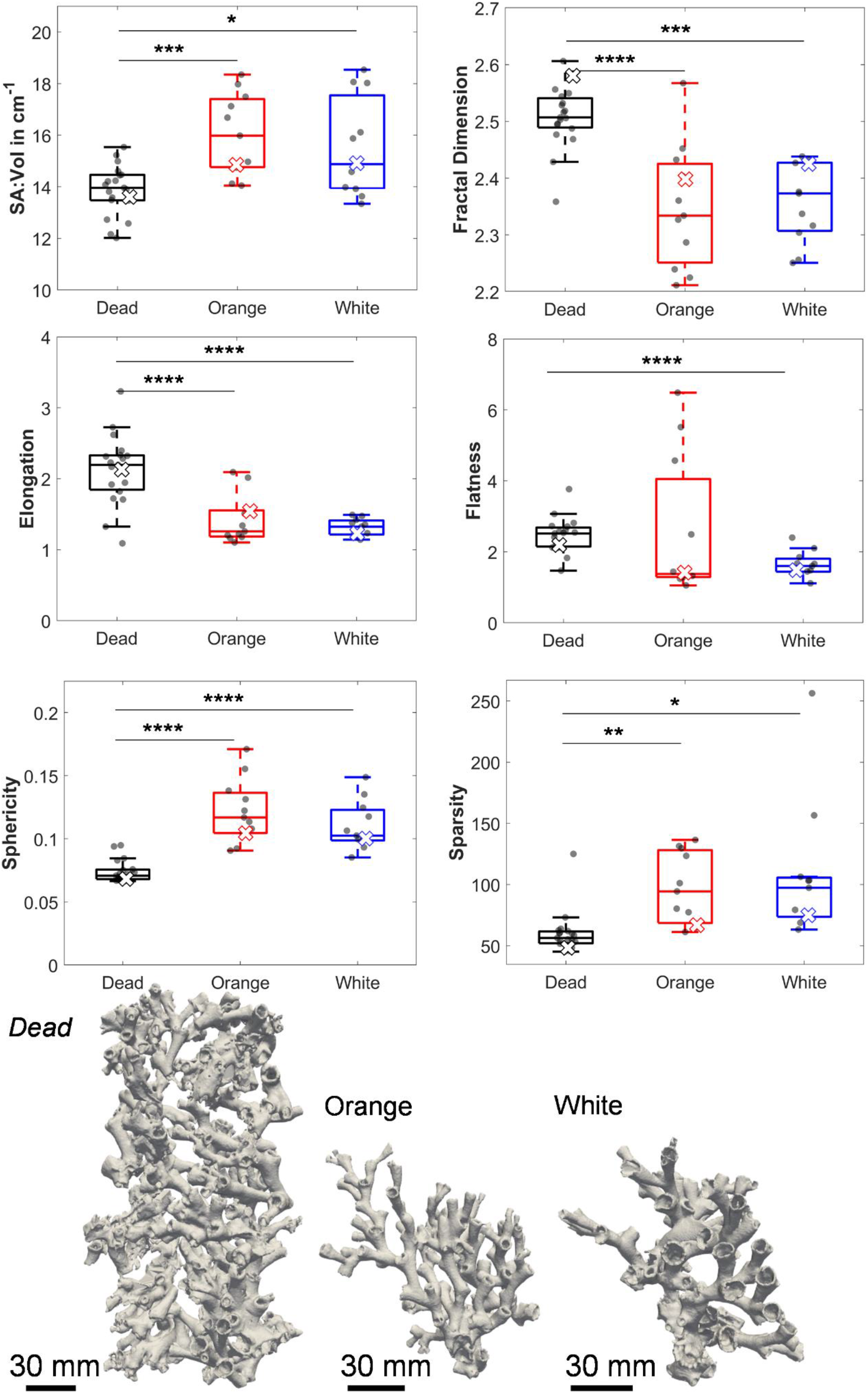
Morphological parameters of *L. pertusa* coral colonies. Boxplots of surface area to volume ratio (SA:Vol), fractal dimensions, elongation, flatness, sphericity and convexity for *dead* coral framework, *live* orange coral and *live* white coral specimens are shown. ‘×’ symbols correspond to shape parameters of the *dead*, orange and white coral specimens shown at the bottom. Significance levels are defined as * p < 0.05, ** p < 0.01, *** p < 0.001, **** p < 0.0001.

### 3.3. Morphology of CWC skeletal branches

Orange corals contained larger branches, with higher volume compared to the white corals, yet wall thickness remained similar (Table 2). *Dead* CWC framework branches were significantly thicker than those from *live* CWC specimens, which resulted in larger area. The larger values of the ellipsoid factor and taper rate for orange corals indicate a more prolate shape and wider opening of their branches. Overall, skeletal branch morphology was highly variable within each specimen (supplementary Figure S2).

**Table 2.** Summary of local morphological parameters. Branch volume (Br.Vol), area (Br.Ar), length (Br.Len), thickness (Br.Th), ellipsoid factor (Br.Ef), and taper rate (Br.Tr) for the pooled data of individual branches of *dead* coral framework, *live* orange coral and *live* white coral specimens are reported as mean (standard deviation) for the n_br_ branches of each group. p-values for the comparison between groups are also included.

|  | Br.Vol<br>in mm<sup>3</sup> | Br.Ar<br>in mm<sup>2</sup> | Br.Len<br>in mm | Br.Th<br>in mm | Br.Ef<br>in - | Br.Tr<br>in mm |
| --- | --- | --- | --- | --- | --- | --- |
| *Dead*<br>($n_{br} = 50581$) | 777.5 (2182.1) | 17.7 (7.1) | 6.1 (3.7) | 2.1 (0.6) | 0.15 (0.31) | 2.2 (2.0) |
| Orange<br>($n_{br} = 1697$) | 1061.2 (2358.3) | 13.7 (7.1) | 7.4 (5.4) | 1.8 (0.4) | 0.25 (0.28) | 2.7 (1.7) |
| White<br>($n_{br} = 3737$) | 640.2 (1427.2) | 11.7 (9.5) | 6.2 (3.9) | 1.9 (0.6) | 0.17 (0.30) | 2.6 (1.9) |
| p-value |  |  |  |  |  |  |
| *Dead/orange* | < 0.0001 | < 0.0001 | < 0.0001 | < 0.0001 | < 0.0001 | < 0.0001 |
| *Dead/white* | < 0.0001 | < 0.0001 | 0.022 | < 0.0001 | 0.013 | < 0.0001 |
| Orange/white | < 0.0001 | < 0.0001 | < 0.0001 | < 0.0001 | < 0.0001 | 0.019 |

Skeletonisation of the CWC structure demonstrated significantly lower number of branches and nodes per unit of skeletal volume of the orange specimens (Figure 8). The higher values for the white colourmorph partially relate to the large number of end branches at distal region of the corallites (Figure 7).

**Figure 7.**
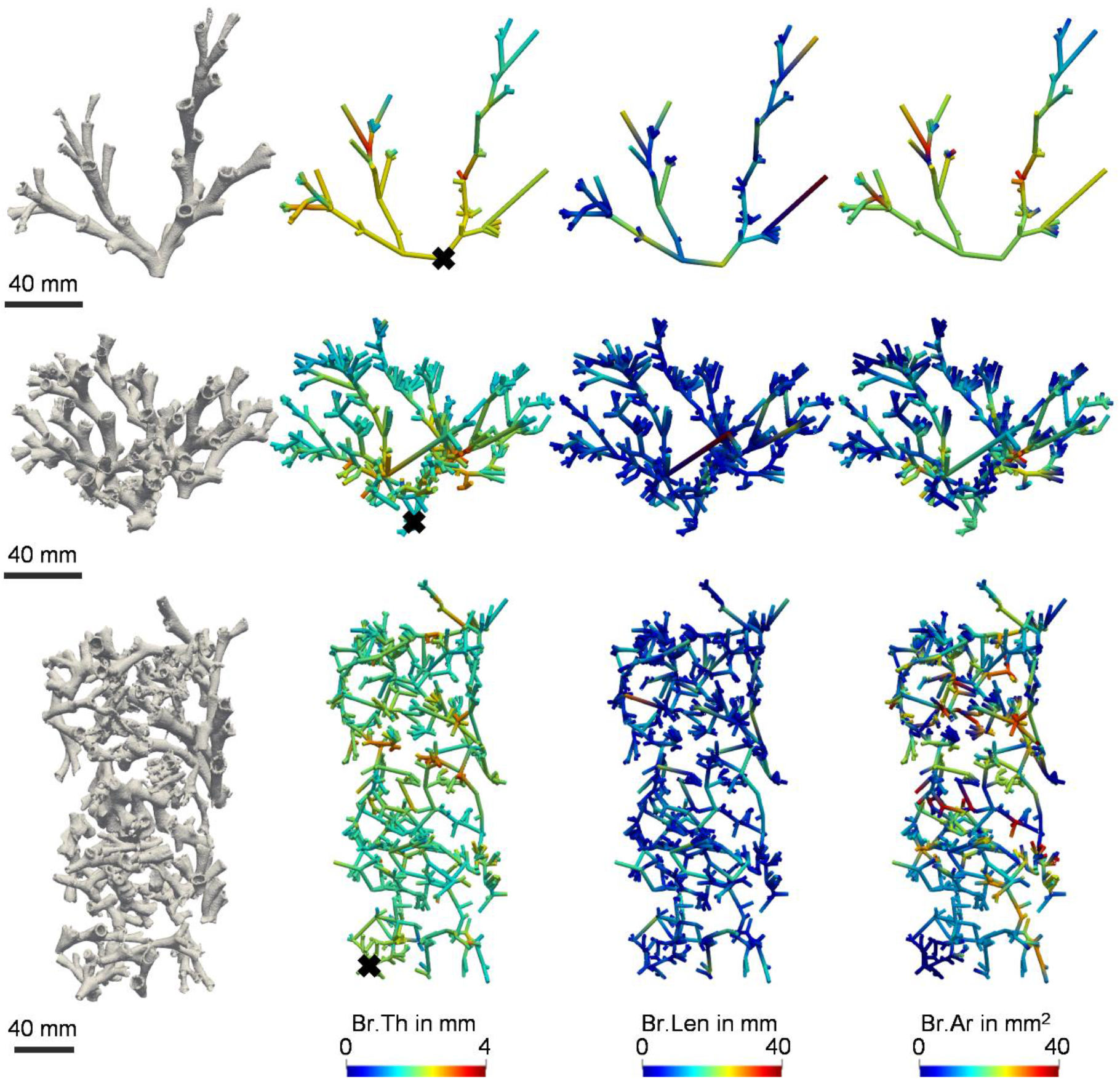
Local morphometry of CWC branches. 3D graphical representation of branch thickness (Br.Th), length (Br.Len), and area (Br.Ar) of the individual branches for three representative *L. pertusa* specimens with increasing levels of complexity: (top) l*ive* orange, (middle) *live* white and (bottom) *dead* framework skeletal structures. ‘×’ symbols correspond to the defined skeletal root (base).

**Figure 8.**
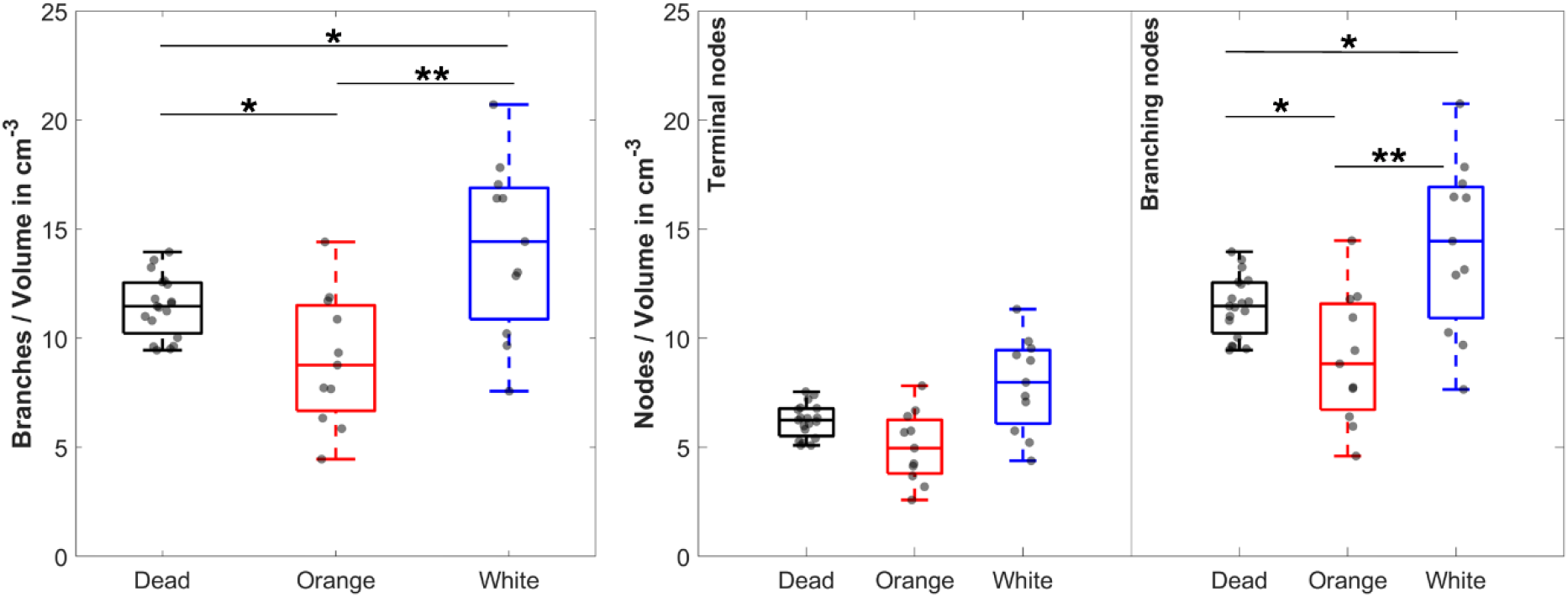
Number of branches and nodes per unit of skeletal volume. Boxplots of the number of branches, terminal nodes, and branching nodes per cm^3^ for *dead* coral framework, *live* orange coral and *live* white coral specimens are shown. Significance levels are defined as * p < 0.05, ** p < 0.01.

### 3.4. Topological descriptors

*B*_*P*_ decreased with increasing radial distance (Figure 9a). Such decrease was faster for the *dead* framework and white specimens. This illustrates that for the orange specimens, newer polyps appear further from the root. Similarly, the distribution of *T*_*BI*_ (Figure 9b) showed a faster decrease for *live* coral specimens, that is, the path to reach the end points of the structure is more direct than for the *dead* CWC framework. The higher complexity of the skeletal structure of *dead* CWC specimens results in increased *τ* (Figure 9c), as well as a larger *V*_*d*_ (Figure 9d), which decreased for the more distant branches.

**Figure 9.**
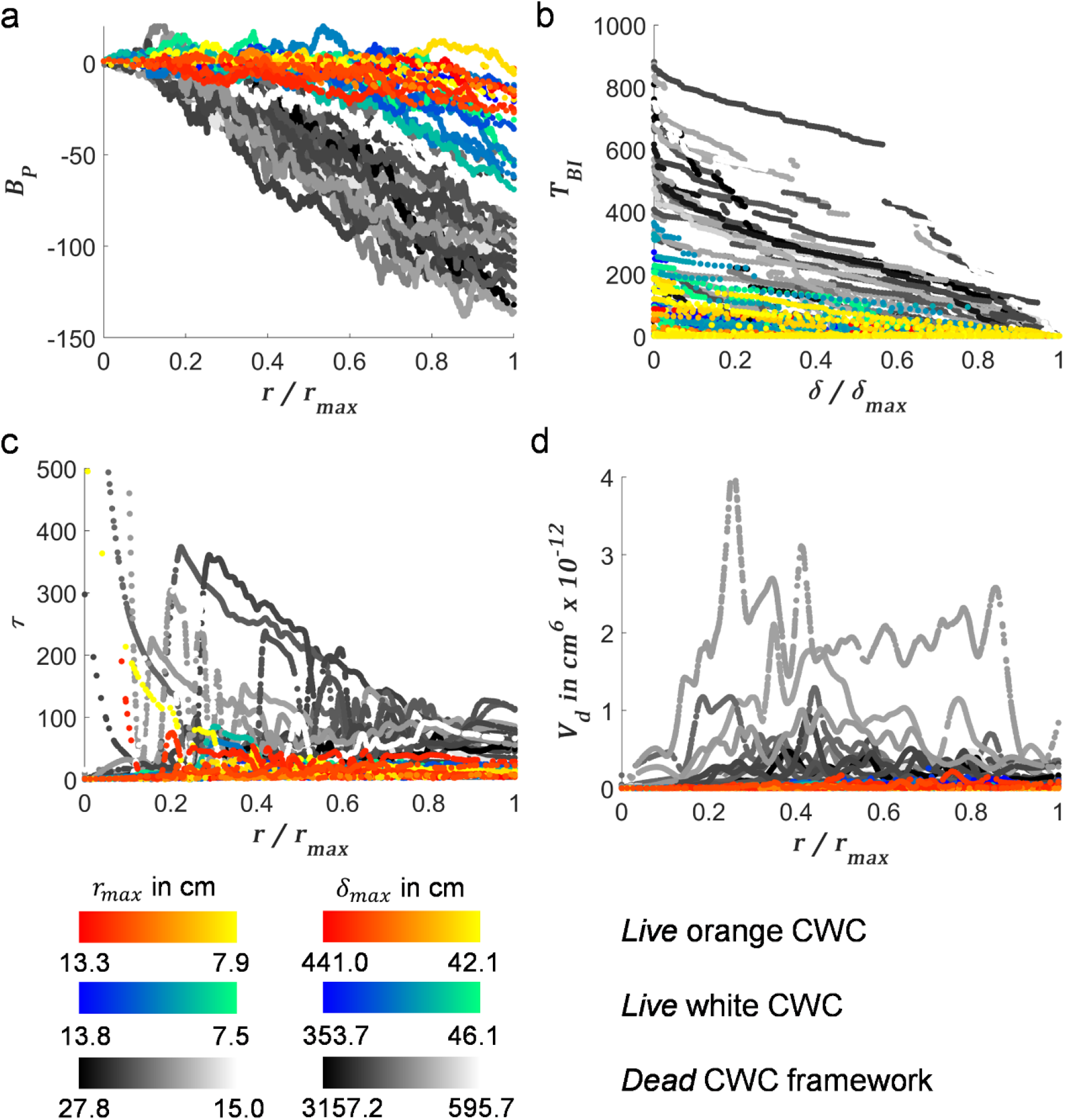
Topological descriptors. (a) Branching pattern (*B*_*P*_), (b) terminal branch index (*T*_*BI*_), (c) tortuosity (*τ*), and (d) volume distribution (*V*_*d*_) as a function of the radial, *r*, or path distance, *δ*, from the skeletal root. Scatterplots for each variable are colour-coded for each group and maximum radial, *r*_*max*_, or path distance *δ*_*max*_.

### 3.5. Principal component analysis (PCA)

At the colony level, the first principal component (PC1) of the PCA explained 38.5% of variation (Figure 10a, b). The projection of white CWC specimens in the score plots was largely aligned in the direction of PC1 (Figure 10a), which was primarily contributed by the fractal dimension and the elongation (Figure 10b). The second principal component (PC2) explained 34.5% of variation, with a major contribution of the number of branches and nodes. A clear positive correlation between those variables was identified, as well as a negative correlation between elongation and sparsity. Fractal dimension and elongation, sphericity and sparsity were also positively correlated. Flatness had the lowest contribution on both PCs. At the skeletal branch level, PC1 explained 76.1% of the variation, whereas the PC2 explained only 11.1% of the variation (Figure 10c, d). *B*_*P*_ and Br.Tr were positively correlated and largely aligned with PC2, while showing a negative correlation with Br.Th, Br.Len, Br.Ar, *T*_*BI*_ and *τ. Dead* CWC specimens showed the lowest variability of the morphometric parameters at the colony level (Figure 10a), but the highest variation at the skeletal branch level (Figure 10c).

**Figure 10.**
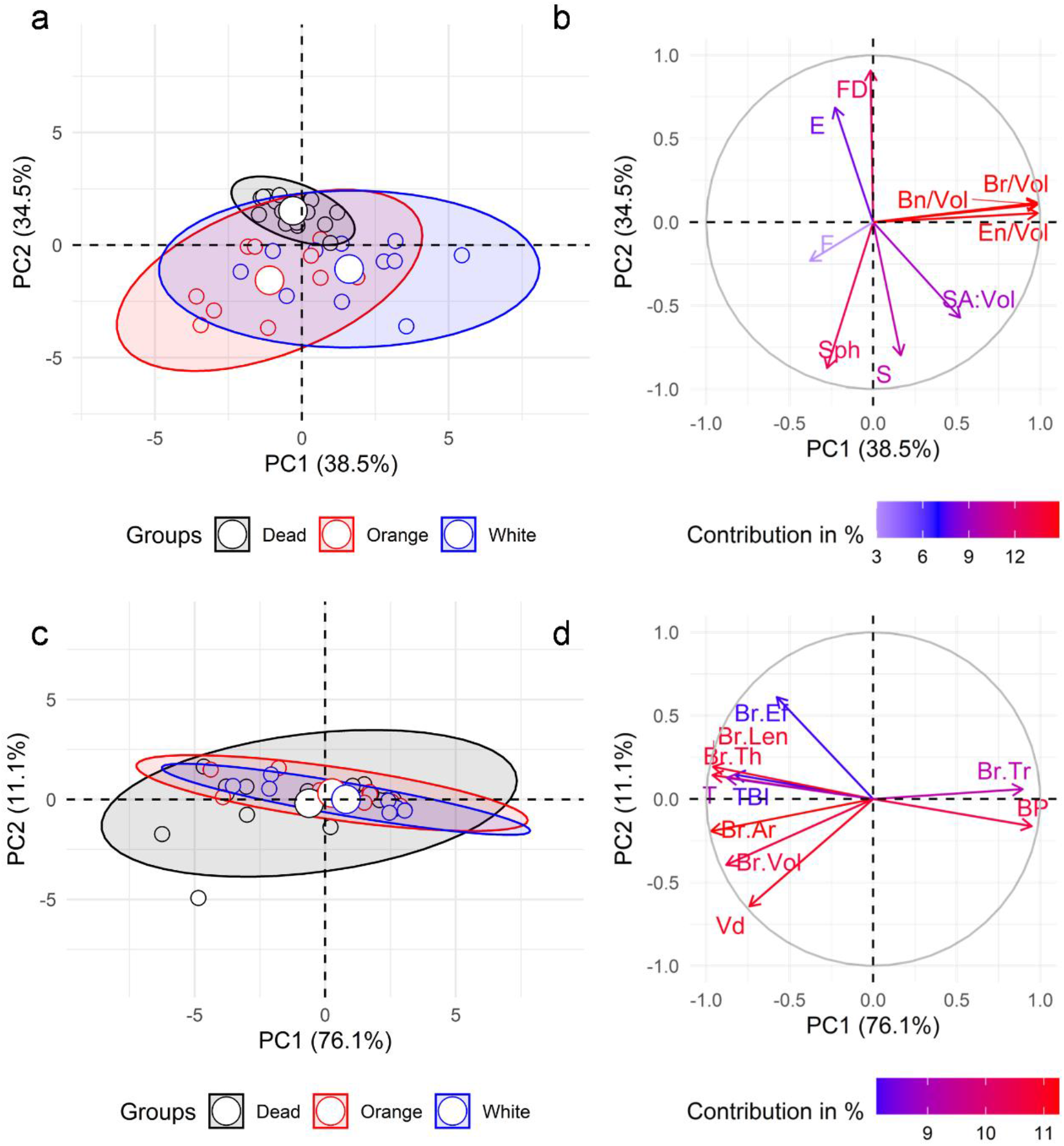
Principal component analysis. (a, c) Score and (b, d) loading plots resulting from PCA for the morphological analysis at (a, b) colony and (c, d) skeletal branch level. Points in (a, c) represent the projection of the analysed CWC specimens. Arrows in (b, d) indicate variable loadings. *S*_*ph*_: sphericity; *S*: sparsity; *SA*: *Vol*: surface area to volume ratio; *F*_*D*_: fractal dimension; *E*: elongation; *F* flatness; *Br/Vol*: number of branches; *B*_*n*_*/Vol*: branching nodes; *E*_*n*_*/Vol*: ending nodes; *B*_*P*_: branching pattern; *T*_*BI*_ : terminal branch index; *V*_*d*_ : volume distribution; *τ*: tortuosity; *Br Vol*: branch volume; *Br*. *Ar*: branch area; *Br*. *Len*: branch length; *Br*. *Th*: branch thickness; *Br*. *Ef*: branch ellipsoid factor; *Br*. *Tr*: branch taper rate.

## 4. Discussion

### 4.1. Limitations of the continuum assumption in *Lophelia* pertusa skeletons

We investigated the critical size of coral colonies that allows us to use a mechanical homogenisation approach to study the mechanical vulnerability of CWCs to ocean acidification. We showed that the orthotropic approximation of the stifness tensor for CWC skeletal structure converges at ∼13 cm edge length, which reflects the critical size for the continuum assumption to hold for *L. pertusa* skeletons at the structural level. This critical size corresponds to 5 to 6 times the mean spacing between skeletal branches. Similar results are seen in trabecular bone mechanical studies in which average continuum properties are sufficient when considering volume elements over 5 intertrabecular lengths [11]. For the two specimens under study, such critical size may include >800 branches. Only two specimens were investigated, which is usually not sufficient to provide a representation of the elastic properties of CWC skeletal structure. This was limited by the availability of 3D data on large CWC colonies, and it explains the motivation behind using mirrored specimens. The consistency in the statistical evaluation of the individual branches, however, indicates that the determined critical sizes are robust and applicable to other CWC samples. In addition to this, we demonstrate applicability of the methodology so that our results can be easily confirmed if larger sample become accessible.

### 4.2. Morphological variability of *L. pertusa* skeletal structure

We used six shape variables to describe how CWC colonies occupy the space. Variations in volume compactness captures a gradient from *dead* to *live* CWC specimens (Figure 10). This is indicative of the open branching structure of *live* colonies, which is optimised for food particle capture, opposite to the more planar structure of *dead* framework that does not require access to food. Moreover, the ability for building reef frameworks increases for colonies with high compactness [38]. However, compactness constrained surface complexity, as previously shown by Zawada et al. [33]. Thus, *live* colonies with higher levels of compactness tended to be smooth (i.e., higher SA:Vol and lower fractal dimension). The increased SA:Vol of *live* colonies also implies increased exposure to the environment to capture food. Indeed, variation in surface complexity relates to competition and resource use [33], where colonies with higher structural complexity have less access to those resources (e.g., nutrients), but can have more polyps packed within a given space [39]. Thus, the death of the CWC colonies may result from the inability of their polyps to source prey and nutrients once a threshold in complexity is passed, as increased complexity will directly impact local hydrodynamics and, consequently, growth [40]. From a mechanical perspective, the highly packed structure of the *dead* framework serves to support living colonies by sustaining external loads.

We quantified the morphology of *L. pertusa* skeletal branches based on a skeletonisation of 3D CT images, as first proposed in [34]. While Kruszyńsky et al. [34] focused on branch diameters and spacings, which are controlled by a combination of hydrodynamics and genetics [41], we extended the analysis to other parameters that may influence the load-bearing capacity of *L. pertusa* colonies, such as wall thickness, branch length, cross-sectional area and volume. Some of these parameters (e.g., thickness and length) have only been quantified using linear measurements from small coral fragments [20,26], consequently, restricting the analysis to a small number of corallites. However, *L. pertusa* colonies are made up of hundreds to thousands of corallites. Thus, the method we present here provides an efficient tool to assess intraspecific morphological variations in CWCs. In line with [26] and [20], we showed a large variation in size and shape for the analysed branches. The measured mean taper rate (i.e., difference in diameter between top and bottom of the corallite) aligns well with previous studies [20,26], highlighting that our CT image-based approach is able to capture slight variations of branch morphology. However, we reported thicker wall values as we consider the mean thickness of the entire branch as opposite to the thinner wall at the top of the corallites in [20,26]. Accurate measurements of the wall thickness in CWC skeletons is important as ocean acidification induces dissolution of the skeletal wall material, decreasing the load bearing capacity of the entire structure [3–5].

In addition to the branch size and shape, we analysed four topological descriptors that describe the branching characteristics of CWC structures. These descriptors confirmed the higher complexity of the *dead* framework, as seen in the increased tortuosity and volume distribution, which may be a consequence of dense packing of several colony fragments. Although the increased volume distributions (i.e., higher mass density) may represent an advantage of the *dead* framework to support living colonies under mechanical stressors, the bare *dead* coral skeleton, which lacks protection by organic tissue or defence mechanisms [42], is more vulnerable to dissolution [4] and bioerosion [27,43], and consequently, mechanical damage [44]. Therefore, degradation of skeletal branches in the *dead* framework may compromise the stability of the entire colony, leading to a rapid collapse and, consequently, loss of biodiversity support [4].

Our principal component analysis did not point to clear branching pattern-specific differences in branch morphology between *dead* and *live* skeletons, or between colourmorphs (Figure 8). This suggests that we cannot simply skip morphological features when developing mechanical surrogate models.

### 4.3. Towards reef-scale modelling of cold-water corals

The investigation of timescales for CWC reef crumbling relies on the development of computational tools that are able to provide accurate and efficient predictions of the mechanical behaviour based on the time CWCs are exposed to acidified waters [4,5]. These tools are currently non-existing, partially due to the lack of information on the structure-function relationships of CWCs at the structural level. Here, we showed that homogenised FE models of coral colonies may be used to estimate the risk of crumbling of CWC colonies when these surpass the identified critical size of 5 interbranch lengths. Similarly to previous studies in other multiscale structures [45–47], the stiffness and strength of the individual finite elements can account for the skeletal volume fraction and architectural information (Figure 11a). This approach outperforms current models that considered a uniform coral skeletal structure [13,14], thus, missing the influence of CWC branching structure on the mechanical response. However, CWC reefs comprise multitude of colonies of varying sizes and shapes, with the size of some of those colonies smaller than the critical size we determined (e.g. on Tisler Reef (Norway) almost 60% of colonies of sizes smaller than 30 cm [23]). In such case, continuum modelling would be unsuitable and must be replaced by explicit models of the branching structure of *L. pertusa* skeletons.

**Figure 11.**
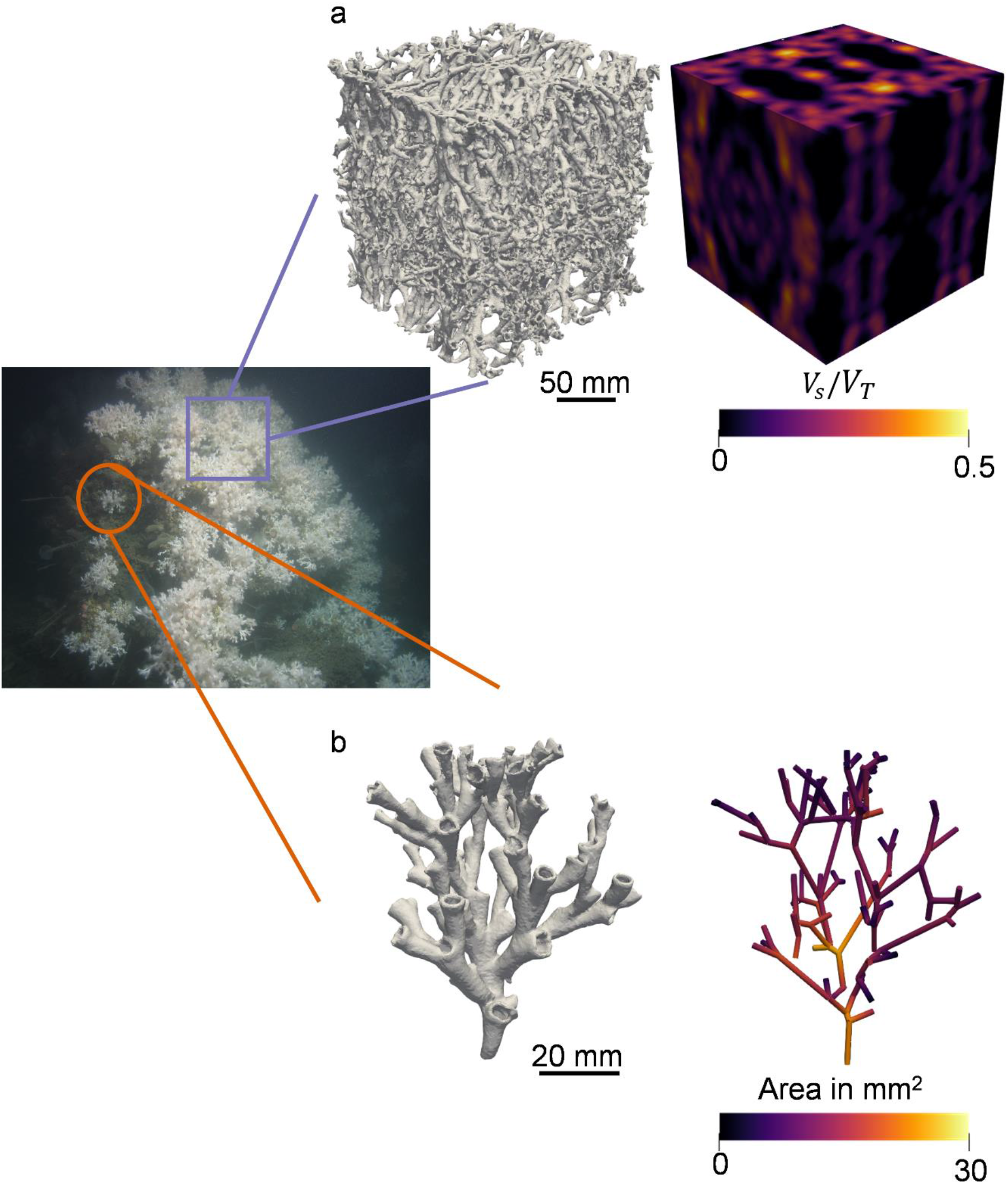
Realms of surrogate models for *L. pertusa* CWC skeletal structure. (a) Large coral colonies (> 13 cm, 5 interbranch lengths or 800 branches) may be modelled using homogenised finite element models based on density (i.e., skeletal volume fraction, *V*_*S*_*/V*_*T*_) and/or fabric information. (b) The morphology of the skeletal branches needs to be considered modelling smaller coral colonies, using, for example, nonlinear beams.

The 3D graph representation we presented here may serve to define surrogate models of these smaller colonies consisting of nonlinear beams (Figure 11b). Beam FE models have demonstrated to predict the mechanical properties of complex structures in an accurate and computational efficient manner [48,49]. As at this small length scale *L. pertusa* skeletal structure is not random, the properties of each beam (i.e., skeletal branch) may account for the statistical information of the morphological analysis. Future analyses would need to validate the use of such specimen-specific beam FE models against image-based FE models to confirm its applicability for predictions of the risk of crumbling of CWC skeletal structures.

The expected reduced computational cost of the models we identified here can ease the translation from the mechanical response at the structural level to entire reefs, facilitating surrogate models at the reef scale to investigate the mechanical vulnerability of CWC reefs to ocean acidification. Most importantly, we could model the increased dissolution in the *dead* framework observed when increasing aragonite concentration [4] as a decrease in the skeletal volume fraction in our homogenised models or a reduced branch thickness in our beam surrogates.

Although we provide a detailed morphological evaluation of a large number of *L. pertusa* skeletal fragments, these only represent a fraction of the colony from two unique reefs in Norway. While collection of larger CWC specimens is limited and ethically questionable, future analysis should focus on the characterisation of morphological variations of skeletal fragments collected from different locations. Monitoring reef corals largely relies on 2D measurements of colony size or colony cover [23,44], thus lacking high-resolution volumetric information. To investigate the risk of crumbling of larger colonies we need to investigate new techniques that allow us to infer 3D volumetric information from planar data [50]. Probably the most crucial gap is the lack of information on time estimates for morphological changes due to exposure to ocean acidification. This will require to extend previous mesocosm experiments [4] to different acidification conditions to establish an exposure trajectory of aragonite concentrations and the relationships with ocean acidification induced dissolution. This data would allow us to use the proposed surrogate models as predictive tools to investigate timescales of loadbearing capacity changes on the time CWCs are exposed to acidified waters.

## 5. Conclusion

We show that the mechanical vulnerability of CWC reefs to ocean acidification may be investigated using surrogate models of *L. pertusa* skeletal structure which are computationally feasible. We proposed a size limit that allows the determination of the type of model to use based on the characteristic material architecture of the skeletal structure. Our detailed morphological analysis point to large variations between *L. pertusa* colonies and skeletal branches, as well as *dead* and *live* skeletal structures which need to be considered for the development of rapid monitoring tools for the prediction of crumbling of CWCs. Spatially large CWC colonies may be modelled using homogenised FE models, whereas small colonies may be modelled using specimen-specific beam FE models. Both approaches may allow us to efficiently scale up the analysis to entire reef systems to investigate reef crumbling due to the time CWCs are exposed to acidified waters. Ultimately, this will support future conservation and management efforts by indicating which marine ecosystems are at greatest risk, when they will be at risk, and how much of an impact this will have on the biodiversity they support.

## Acknowledgements

We would like to thank Dr Carola Daniel for her assistance during CT imaging and captains and crews of research cruises POS455 (2013) and POS473 (2014) with RV POSEIDON during which Norwegian samples were collected.

