## supplementary material for "Towards modelling cold-water coral reef-scale crumbling: Including morphological variability in mechanical surrogate models"

### Supplementary information

#### S1. Image post-processing

We used Python 3.8 libraries SimpleITK and scikit-image for image post processing. First, we resampled the images to an isotropic voxel size of 0.351 $mm^3$ for the Norwegian specimens, 0.662 $mm^3$ for the Rockall Bank specimen, and 0.445 $mm^3$ for the West of Shetland specimen. We reduced noise using a recursive Gaussian filter with a filter width of $\sigma = 0.3$. Thereafter, we segmented coral skeletons and cavities individually. We segmented the coral skeletons using a maximum entropy algorithm and we used a connected component analysis to remove isolated regions that were not attached to the skeleton (Figure 2a, d). We then segmented cavities within the skeleton in three steps. First, we applied a 3D morphological closing filter with a kernel radius of 6 pixels followed by a 3D binary dilation with a kernel radius of 2 pixels to the segmented coral skeleton to create a coarse mask contour image of the combined coral skeleton and cavities. This mask was then refined using an iterative 3D geodesic active contour (3D-GAC) algorithm (Ohs et al., 2021), which allowed us to identify the external contour of the skeleton (Figure 2b, e). We obtained a binary image containing only the cavities by subtracting the binary coral skeleton from the contour mask image. We labelled individual corallite calices using a hierarchical watershed on the 3D distance map (Figure 2c, f).

To quantify the local size of both the coral skeleton and corallites, we computed 3D thickness maps (Hildebrand et al., 1999) of the contour and cavities mask images, respectively (Figure 2g,h). We assessed the local shape of the coral skeleton by measuring the 3D ellipsoid factor maps (Doube, 2015) on the contour mask image (Figure 2i). Finally, we computed the mean spacing between skeletal branches from the 3D spacing maps of the contour images. Thicknesses, spacing, and ellipsoid factor were computed using BoneJ (Doube et al., 2010) plugin in Fiji (Schindelin et al., 2012).

#### S2. Representative volume element (RVE) of cold-water coral skeletal structure

An RVE can be defined as the smallest volume element of a heterogeneous structure for which a macroscopic constitutive representation is sufficiently accurate to model the mean constitutive response (Drugan and Willis, 1996). Therefore, an appropriate size of an RVE of the skeletal structure of CWCs should be found that consider: (i) a large enough number of heterogeneities to be statistically representative of the corals' structure; (ii) a size small enough so that it can still be considered as a material point from a macroscopic point of view. The RVE domain shall comply with the Hill condition (Hill, 1963), which states the necessary and sufficient conditions for equivalence between energetically and mechanically defined properties of elastic materials:

$$\langle \boldsymbol{\sigma} : \boldsymbol{\varepsilon} \rangle = \langle \boldsymbol{\sigma} \rangle : \langle \boldsymbol{\varepsilon} \rangle \qquad (1)$$

This means that the average of the product of the stress $\boldsymbol{\sigma}$ and strain $\boldsymbol{\varepsilon}$ tensors (microscale) equals the product of their averages. Similar to trabecular bone (Pahr and Zysset, 2008), we do not strive to find an RVE where (1) is exactly fulfilled but where we can obtain a usable approximation.

Numerical techniques, such as the FE method, can be used to approximate the critical size for an RVE by analysing the size dependence of the elastic symmetries and properties of the structure (Kanit et al., 2003). These properties can be estimated from the stiffness tensor, $\mathbb{S}$, using a direct mechanics approach through an optimisation procedure where the best orthotropic representation of the stiffness tensor may be found (van Rietbergen et al., 1995). Here, we approximate the critical size of an RVE for CWC skeletal structures by analysing the convergence of the orthotropy assumption.

### S3. Orthotropic stiffness tensor components

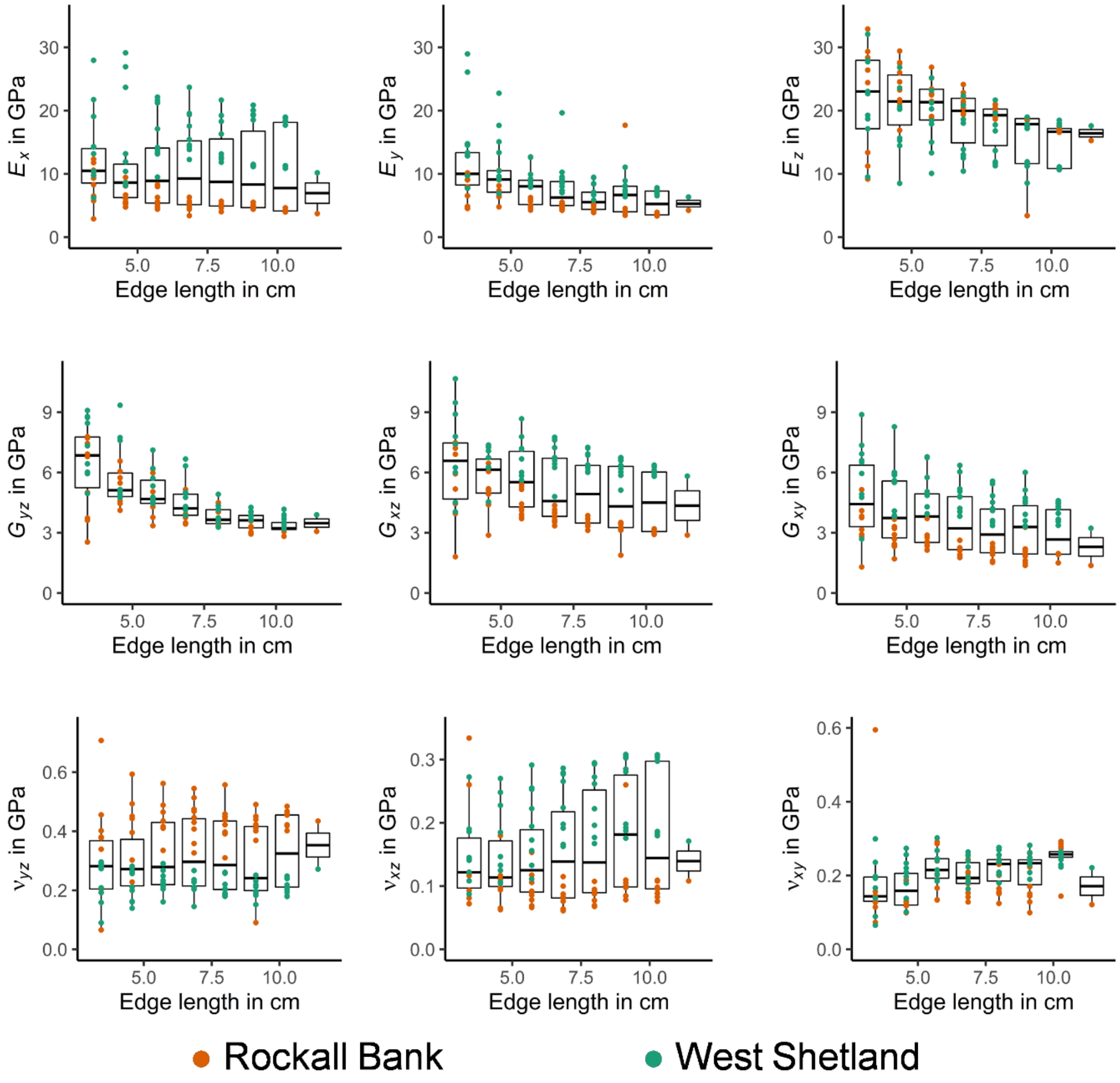


**Figure S1. Orthotropic stiffness tensor components**. The components of the orthotropic stiffness tensor, $\mathbb{S}^{ORT}$, computed on volume elements of varying sizes virtually extracted from two representative *L. pertusa* specimens from Rockall Bank and West Shetland (Figure 3). The convergence of the Young's ($E_1$, $E_2$, $E_3$) and shear ($G_{23}$, $G_{13}$, $G_{12}$) modulus with increasing edge lengths is shown. Poisson's ratio ($\nu_{23}$, $\nu_{13}$, $\nu_{12}$) was minimally influenced by the edge length of the volume elements.

### S4. Morphological parameter of CWC skeletal branches

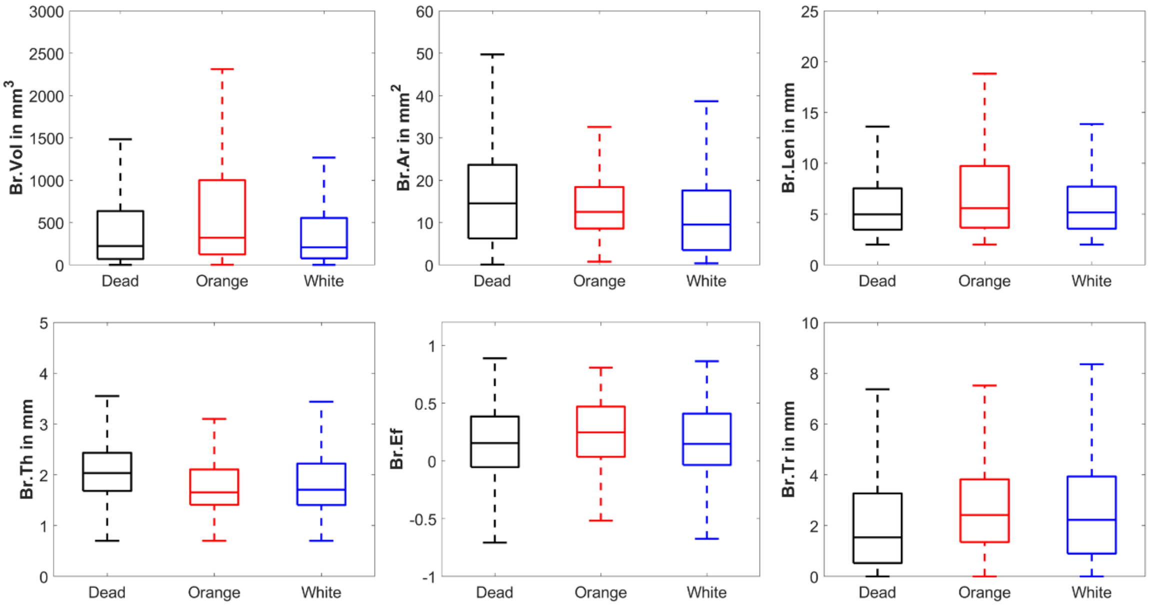


**Figure S2. Morphological parameters of *L. pertusa* skeletal branches.** Boxplots of branch volume (Br.Vol), branch area (Br.Ar), branch length (Br.Len), branch thickness (Br.Th), branch ellipsoid factor (Br.Ef), and branch taper rate (Br.Tr) for *dead* coral framework, *live* orange coral and *live* white coral specimens are shown.
